# Human retinal organoid single-cell atlas allows to reconstruct retinal development at high resolution and identify nature restricted transcriptional states in vitro

**DOI:** 10.1101/2025.08.02.668295

**Authors:** Emil Kriukov, Everett Labrecque, Nasrin Refaian, Petr Baranov

**Affiliations:** Massachusetts Eye and Ear, Boston, MA; Department of Ophthalmology, Harvard Medical School, Boston, MA

## Abstract

The vertebrate retina is a highly specialized neural structure crucial for the initial capture and processing of visual information. Perturbations in retinal development contribute significantly to congenital and acquired forms of blindness, underscoring the necessity of elucidating human retinal ontogeny. Human pluripotent stem cell-derived retinal organoids have emerged as a powerful model capable of self-organizing and recapitulating key aspects of retinal morphogenesis, offering valuable insights into retinal development and disease. Despite their utility, the extensive cellular heterogeneity, dynamic state transitions, and existence of unique, the development of this system has not yet been fully resolved, posing significant challenges for interpreting their developmental fidelity and therapeutic applicability.

Here, we present the Human Retinal Organoid Cell Atlas (HROCA), a comprehensive single-cell transcriptomic reference spanning retinal organoid development from Day 10 to Day 365, integrating data from 15 studies, 71 batches, and encompassing 458,309 cells. HROCA provides a detailed benchmark for organoid differentiation, identifying known retinal cell classes and novel cellular states at single-cell resolution. By associating HROCA with the human fetal retina atlas (HFR), we offer an in-depth comparison between in vitro and in vivo retinal development, revealing protocol-specific differences, limitations in neuronal maturation, and developmental timing discrepancies. HROCA expands our understanding of retinal developmental plasticity, highlighting unique cell populations such as chimeric, non-canonical, and apatride cells, which do not conform to classical retinal cell types but exist consistently across developmental timelines in vitro. This atlas serves as a crucial resource for understanding retinal development, evaluating organoid models, and guiding future improvements in regenerative medicine strategies.

## Introduction

The vertebrate retina is a highly specialized neural tissue responsible for capturing light and initiating visual processing. Its complex, layered architecture arises through a tightly orchestrated sequence of developmental events, giving rise to diverse neuronal and glial cell types with precise spatial and temporal organization.[1] The disruptions to retinal development underlie many congenital and acquired blinding diseases, making the elucidation of human retinal ontogeny crucial for both basic biology and regenerative medicine.

Three-dimensional human retinal organoids were one of the first systems that demonstrated the ability of pluripotent stem cells to self-organize and undergo lineage specification, followed by recapitulation of the development.[2] These self-organizing structures recapitulate many aspects of retinal morphogenesis, providing unprecedented access to developing human retinal tissue.[3] This platform enabled the studies of retinal development and disease modeling, followed by organoids application in drug discovery and as a substrate in cell replacement. However, this model’s cellular heterogeneity and dynamic transitions have not been comprehensively resolved, limiting our understanding of their fidelity to in vivo development and their limitations for clinical applications.

Also, there is a lot of knowledge accumulated on in vivo retinal development and the description of possible developmental cell states, i.e. “nature restricted” states. This has not been the case for retinal organoids, and their heterogeneity and plasticity suggest there is more of possible developmental states in vitro than it is known in vivo.

Here we present a comprehensive single-cell atlas spanning human stem-cell-derived retinal organoid development, human retinal cell organoid atlas (HROCA), integrating 15 different studies/protocols and 71 individual batches, with a total of 458.309 cells from Day 10 to Day 365 of organoid development. This atlas provides a reference framework to benchmark organoid differentiation, reveals novel cellular states, and describes the known cell classes at a single-cell resolution. We also provide the detailed comparison of in vitro and in vivo retinal cell class development through association of HROCA with the human fetal retina atlas (HFR). It allows to explore timepoint/protocol-specific differences, the limitations of the neuron maturation in vitro, and correlate in vitro and in vivo timelines of major biological functions. We demonstrate how HROCA expands our understanding of “natural-liimited” developmental patterns by bringing chimeric, non-canonical and apatride populations into the bigger picture of possibly existing states.

## Methods

### Single-cell RNA sequencing data obtaining and processing

Public scRNAseq data from healthy WT human retinal organoids[4–19] were collected from the GEO repository using fastq-dump from SRA-toolkit version 3.1.1 for the .fastq and .bam files. .bam files were converted to .fastq.gz using cellranger bamtofastq. Using Cell ranger version 8.0.1, we performed cellranger count with the refdata-gex-GRCh38-2024-A reference genome.

Single-cell RNA sequencing data analysis was performed in RStudio v 4.4.1 using filtered_feature_bc_matrix provided in the outs folder created by cellranger count. The filtered Cell ranger output data was processed independently following the Seurat pipeline, using Seurat v. 4.4.0[20] . The pipeline includes modified ScaleData() function, with the argument vars.to.regress = c(’percent.mt’,“percent.rb”,“S.Score”,“G2M.Score”).

### Data Integration and Preliminary Atlas Generation

To integrate the datasets upon independent processing of every object, we used scVI v. 1.3.2[21] . First, we perform hyperparameter fine-tuning with the following choice of parameters: n_layers = [2, 3, 4, 5], n_latent = [20, 30, 40, 50], n_hidden = [64, 128, 256], gene_likelihood = “zinb”, lr = [1e-4, 1e-2]. Following these parameters for training, we select the top-5% performing models based on the validation_loss metric. The selected models are further used in benchmarking to assess the “Batch correction”, “Bio conservation”, and “Total” parameters through scIB benchmarking[22] . Based on these scores, we select the best performing model for the integration. These metrics are available in **Figure S1** and **Supplementary Table 1** for validation loss, and **Figure S2** for scIB benchmarking.

We used the model with the following parameters to perform integration: n_layers=2, n_latent=30, max_epochs=100, n_hidden=256, early_stopping_patience=5.

### Protocols metadata building, assessment, and unification

For each of the GSE datasets and GSM samples obtained, we collect the metadata that includes both dataset-specific (timepoint/age, chemistry, assay, etc.) and protocol-specific (stem cell origin, plate type, attachment, each media/factor/supplement, their duration, and concentration, where applicable). Due to the individual approach in protocol description, we focused on preserving the metadata relevant for the organoid, but not stem cell culture. We harmonized and unified the available metadata (Pyruvate and Sodium Pyruvate to Pyruvate, mTeSR1 and mTeSR Plus to mTeSR, retinoic acid and all-trans retinoic acid to retinoic acid, etc.), however, we maintained the original details where possible, avoiding harmonization that leads to metadata loss. The complete protocol-specific metadata is available as **Supplementary Table 2**.

### Atlas Annotation

On the scVI-based UMAP embedding, we resolve the clusters using Leiden[23] algorithm (resolution = 2) and perform cluster annotation using both canonical or known markers together with cluster-specific differentially expressed genes from scanpy[24] and NSForest[25] . We utilize pan-human-azimuth to explore potential predictions from the model. Upon manual annotation, we preserve the annotation overlapping between the manual and pan-human-azimuth[20] approaches in single-cell manner. In this manner, we were able to annotate cycling RPC, amacrine, horizontal, part of rod and cone PhR populations.

For the rest of the populations, we annotate them using manual approach, and once 90% of the cells had the annotation resolved, we pass the .h5ad atlas, scVI model, and annotation metadata into scANVI[26], preserving the “Unknown” annotation label for the clusters we were unable to resolve.

Following this, with the new post-scANVI embedding and clustering, we transfer the annotation for the whole cluster from single cells, if at least 95% of single cell within this cluster have the same identity. For the rest of the clusters (7 clusters we could not assign the annotation to), we repeat the iteration with DEGs search, and finalize the annotation.

The fully annotated atlas was passed to scANVI for the final scANVI-based UMAP embedding.

### CytoTRACE2 and Mellon

Upon the annotation, the resulting .h5ad object was converted to .h5Seurat using SeuratDisk v. 0.0.0.9020. To avoid the issues related to objects structure inconsistency when transferring it from Python Anndata to R S5 object structure, we loaded only the object matrix, and wrapped it with metadata and embeddings further on in a manual manner. Mellon[27] pipeline was performed using default settings using Mellon v. 1.6.1. For CytoTRACE2 v. 1.1.0[28] we used Seurat object as input, and cytotrace2() was called on the counts slot, with batch size of 10000 cells, smooth batch size of 1000 cells, models and smoothing parallelization on. To avoid further issues with conversion from .h5Seurat to .h5ad, CytoTRACE2-related object@meta.data was saved as .csv file and added to the initial .h5ad object.

### scPoli timepoint transfer from HFR v.1

To perform timepoint transfer, we used previously described human fetal retina atlas[29] (HFR) as a reference atlas. HROCA timepoints were binned in a functional/batch-sufficient manner, and scPoli[30,31] model was created using 20 embedding dimensions, 30 latent dimensions, and zero-inflated negative binomial as a parameter for reconstruction loss. Layers architecture was adjusted to [256]*2 as found to be the best performing for scVI. Both model and query were trained using 50 n_epoch, with 40 being pretraining epochs. Unlabeled prototype training setting was set to False.

### The parsing and analysis of cell culture protocols

We perform hierarchical dendrogram using vegan v. 2.6-4 to represent the protocol composition within the main protocol metadata columns that include stem cell origin, attachment, plate type, and number of cells per well. For the other metadata in the protocols, that includes medias and supplements, we harmonize the metadata and demonstrate the heatmap of factors present in every protocol.

Furthermore, to access the correlation between the protocol and cell class, we perform RDA. For each of the protocols the raw number of cells per cell class were extracted, and the frequency was calculated within each protocol.

We further split the protocols into protocols and timepoints, where available. Following the same frequency-based cell composition approach, we build UMAP using 5 dimensions for dimensionality reduction, to plot every protocol_timepoint observation on the embedding.

### ssGSEA pathway analysis

ssGSEA pathway analysis was performed using escape v. 2.5.0[32] package on the Hallmark and C5 selected pathways from Molecular Signatures Database. These particularly include “GOBP_NECROPTOTIC_PROCESS”, “GOBP_PROGRAMMED_NECROTIC_CELL_DEATH”, “GOBP_OPTIC_NERVE_MORPHOGENESIS”, “GOBP_NEURON_FATE_COMMITMENT”, “GOBP_STEM_CELL_PROLIFERATION”, “HALLMARK_APOPTOSIS”. Upon the analysis, normalized enrichment score was min-maxed from 0 to 1 and presented as a heatmap for both HROCA and HFR.

### Jaccard similarity

For both the HROCA and HFR, we identified cell class–specific marker genes using the scanpy implementation of the Wilcoxon rank-sum test (sc.tl.rank_genes_groups). The analysis was performed independently for each atlas, yielding ranked lists of differentially expressed genes (DEGs) for each annotated cell class. For each cell class in both atlases, we selected the top 10/100/300/1000/3000 ranked marker genes based on the Wilcoxon test statistics. These gene sets were extracted from the resulting marker tables. Genes were grouped by cell class, and only the top N marker genes were retained per type for downstream similarity analysis.

To quantify the transcriptional similarity between cell types in the two atlases, we calculated the Jaccard index for each pairwise combination of HROCA and HFR cell classes. For each cell class, the top marker genes were converted into sets. The Jaccard similarity was computed as the size of the intersection divided by the size of the union of the gene sets for each pair of cell types, i.e. the classical Jaccard similarity index[33–35] formula.

The resulting Jaccard index values were stored in a matrix with organoid cell types as rows and fetal retina cell types as columns and visualized as a heatmap. The heatmap displays the degree of overlap between marker gene sets, with higher Jaccard values indicating greater transcriptional similarity between cell classes across the two atlases.

## Results

### Human retinal organoid cell atlas provides a comprehensive transcriptional map of retinal development in vitro

We sourced, reanalyzed, and integrated existing human “healthy” retinal organoids single-cell RNA sequencing data to establish a human retinal organoid cell atlas (HROCA) (**Figure 1A**). The resulting HROCA atlas, built of 14 GSE/studies[4–19], 71 individual batches, contains 458,309 cells from 15 retinal organoid differentiation protocols, spanning the timeline from Day 10 to Day 365. We focused on preserving and unifying the original metadata where possible. Cluster identification and annotation was performed using pan-human-azimuth anchoring, and differentially expressed genes (DEGs) together with canonical cell class specific genes on the scANVI-corrected matrix. Upon cluster annotation, we define the following populations of retinal and non-retinal origin: Pluripotent, Cycling Retinal Progenitor, Retinal Progenitor (RPC), MSX1+RPC, Müller Glia, AC/HC/RGC-precursor (PAX6+), Amacrine, Horizontal, RGC, Bipolar/PhR-precursor (OTX2+), PhR precursor, Rod precursor, Rod PhR, Cone PhR 1, Cone PhR 2, Chimeric PhR, Bipolar, RPE, Lens, Astroglial Progenitor, Vascular, Immune. The notable non-retinal populations include Non-Retinal Progenitor 1 and 2, Non-retinal Glia, Non-Retinal Neuron, and Epithelium. We also describe the population of Apatride cells, the cluster with viable cells that cannot be identified using tissue-, cell-or state-specific markers (**Figure 1B**, **Figure 1C**).

**Figure 1.**
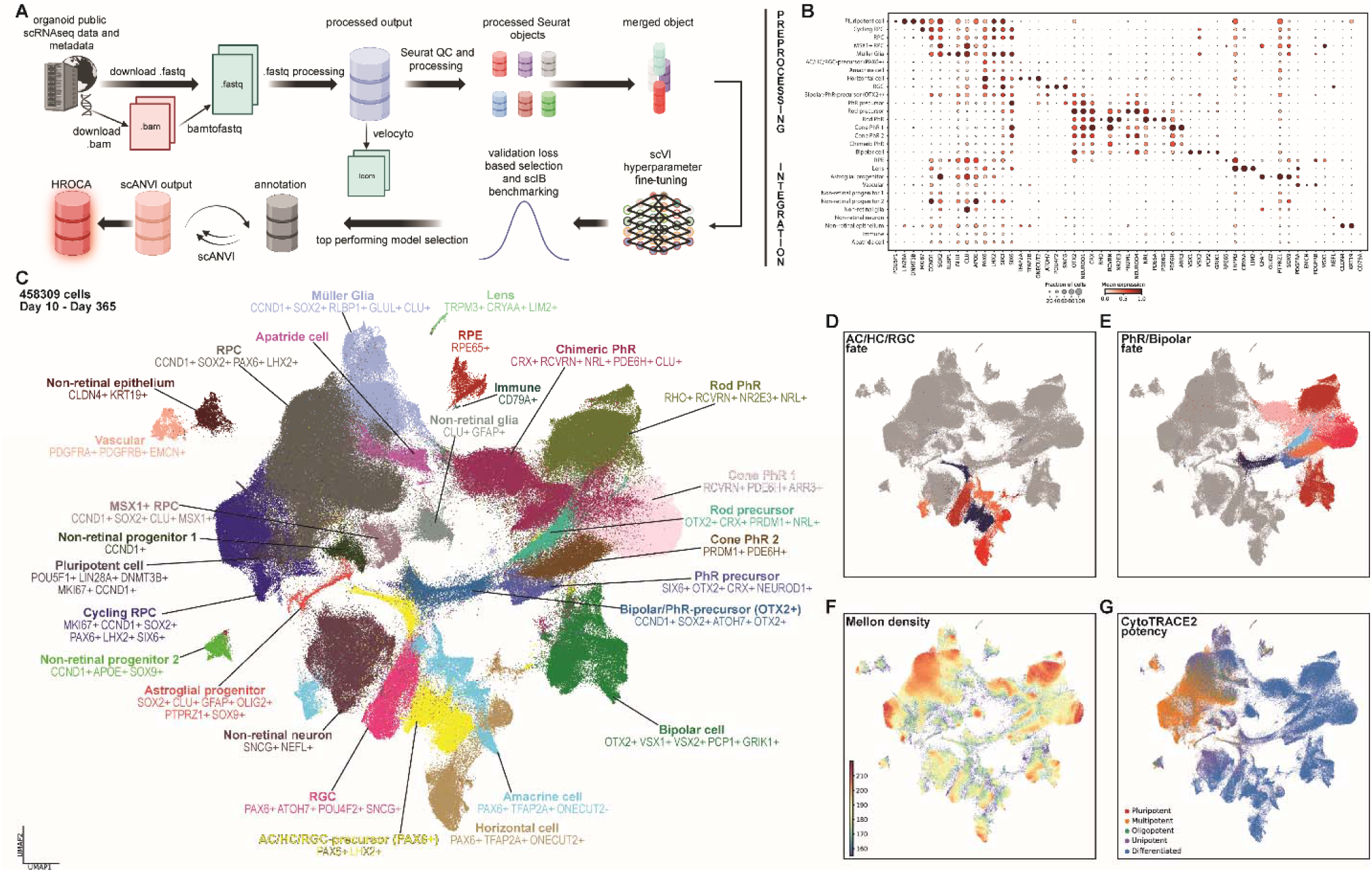
HROCA demonstrates in vitro developing organoid system in single-cell resolution. (A) Pipeline outline, describing the steps from public data parsing and reanalysis, to the final iteration of the atlas integration. (B) Dotplot demonstrating the cell class specific genes for every cell class annotated. (C) Dimplot representing HROCA and its major cell classes present in the dataset. Each of the cell classes annotated is described together with its DEGs. (D) Highlighted subset of AC/HC/RGC fate together with its PAX6+ precursor. (E) Highlighted subset of Bipolar/PhR fate together with its OTX2+ precursor. (F) Featureplot demonstrating cell density parameter achieved by Mellon. Color codes the density, with red meaning high, and blue coding low density. (G) Featureplot representing CytoTRACE2 cell stemness, highlighting the progenitor and fate-committed populations.

Following the annotation of HROCA (**Figure 1C**), we demonstrate two major neuronal branches and their derivatives – PAX6+ precursor of AC/HC/RGC (**Figure 1D**) and PhR/Bipolar from OTX2+ precursor (**Figure 1E**). Together with the application of CytoTRACE (**Figure 1G**) for stemness and potency analysis, and Mellon for cell density and bottlenecks (**Figure 1F**), it allows us to reconstruct the development at higher resolution. The potency of retinal populations resembles normal retinal development with the CytoTRACE2 score high in progenitor populations (RPCs, MSX1+ RPCs), lower in precursor populations (RGC/AC/HC Pax6+, PhR/bipolar OTX2+ and MG precursors) and lowest in more terminally differentiated states. We also observe separate branches of transcriptionally different development trajectories – glial and neuronal. Cell density within the clusters as analyzed by Mellon (**Figure 1F**) is the highest in the progenitor populations (Cycling RPC and RPC), and differentiated cell classes (Müller Glia, Rod PhR, Cone PhR 1, Bipolar, Amacrine, Horizontal, RGC), as well as in Chimeric PhR. Cell density is low in the transitory populations (AC/HC/RGC-precursor (PAX6+), Bipolar/PhR-precursor (OTX2+), Müller Glia to Chimeric PhR “bridge”.

### HROCA allows in silico comparison of variables within multiple organoid differentiation protocols

To further explore the heterogeneity of the organoids integrated in HROCA, we explore 15 retinal organoid differentiation protocols that contributed to the atlas[4–19] . We demonstrate the major HROCA metadata that includes cell class distribution in the atlas, between GSE and timepoints in **Figure 2A**. Additional metadata describing GSE contribution to every cell class and timepoint is available as **Figure S3**. We explore the frequencies of every cell class across the timeline of HROCA (**Figure S4**) and notice the trend, where AC/HC/RGC fate is the most abundant from Day 40 to Day 100, while Bipolar/PhR fate frequency is the highest from Day 80. While early timepoints are characterized with higher frequency of pluripotent and cycling RPC populations, the later ones demonstrate higher differentiated populations frequency, that is noticeable for both retinal (RPE, Bipolar, Müller Glia) and non-retinal or non-canonical (Apatride, Immune, Chimeric PhR) populations. Population size, represented as cell number, for cell class and GSE, and cell class, GSE and timepoint, are available as **Supplementary Table 3** and **Supplementary Table 4**.

**Figure 2.**
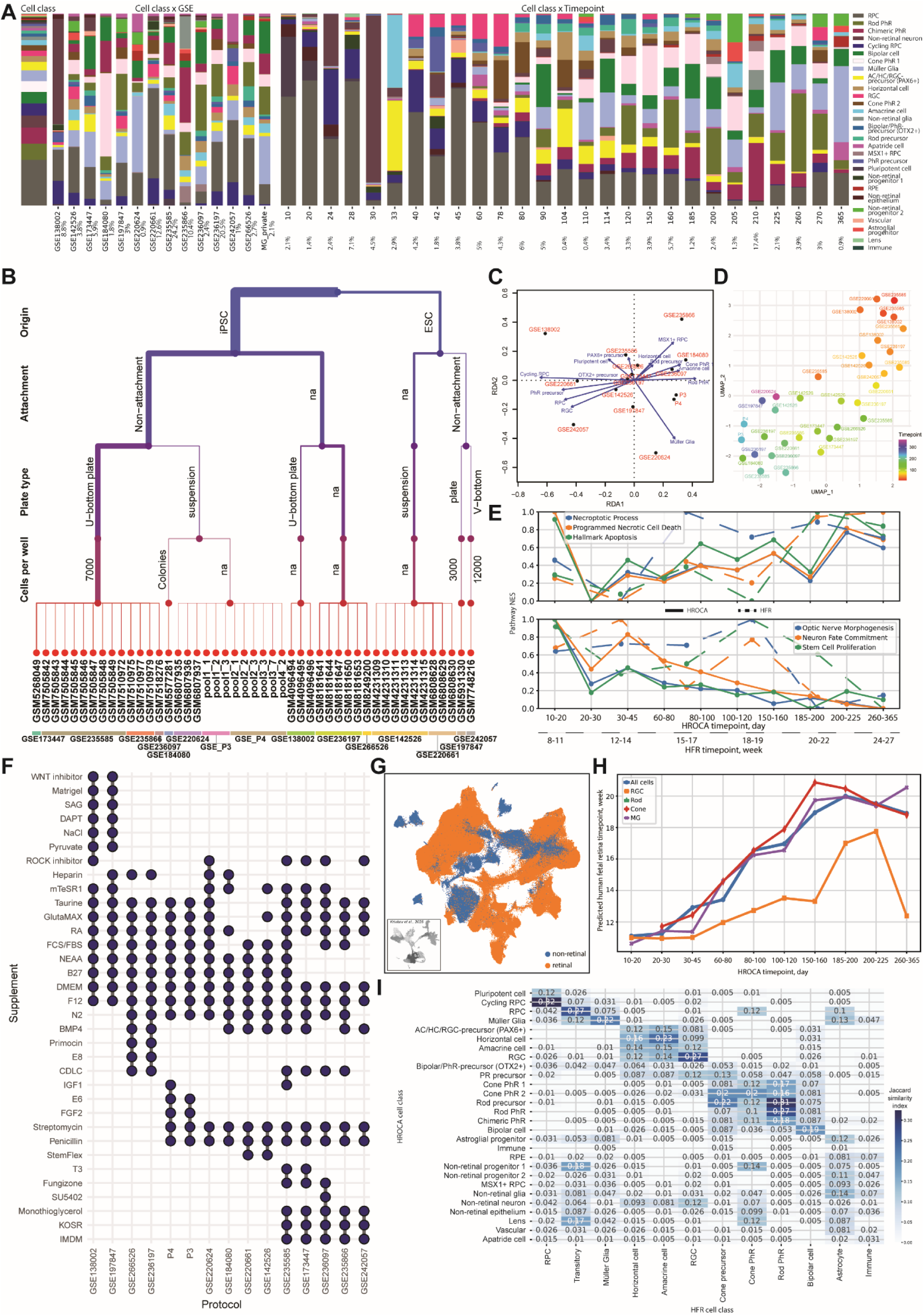
Deciphering HROCA to correlate in vivo and in vitro retinal development. (A) Bar charts demonstrating the distribution of cell classes, cell classes across GSEs, and across the timepoints. Percentage under the labels highlights the contribution of particular GSE/timepoint to HROCA. (B) Dendrogram representing the hierarchical structure of fundamental retinal organoid differentiation protocols parameters across 15 protocols. (C) RDA highlighting compositional prevalence of cell classes across the protocols. Arrows demonstrating the correlation trend between cell class (blue) and protocol (red). (D) UMAP embedding of protocol compositions. Each dot represents a sample that belongs to a protocol and a timepoint. Color codes the timepoint. (E) Chart representing the comparison of selected pathways between HROCA (solid) and HFR (dashed). Chart is separated into two patterns of pathways: increasing (upper) and decreasing (lower) trends. (F) Scatterplot representing the variety of supplements and factors in every protocol. Presence of a supplement/factor is highlighted with a dot. (G) UMAP embedding of HROCA, where color codes the populations present or not in HFR. HFR as a reference atlas is represented in the bottom left part. For the timepoint transfer with scPoli, only the populations present in HFR were taken. (H) Chart demonstrating HROCA timepoints predicted to HFR timepoints for the complete dataset of retinal populations (blue line), RGC (orange line), rod PhR (green line), cone PhR (red line), and Müller Glia (purple line). (I) Heatmap highlighting the similarity between HROCA and HFR cell classes, where color and values on the heatmap represent Jaccard similarity index quantified on cell class DEGs.

Upon hierarchical order of the metadata, we explore the dendrogram of the fundamental factors contributing to the retinal organoids development. These include stem cell origin (hES or iPSC), presence of plate coating, plate type (U-bottom, flat-bottom, suspension), number of cells per well (**Figure 2B**). Furthermore, we explore the protocol-specific factors, such as medias and supplements, defining each protocol (**Figure 2F**). The complete metadata on every factor, its duration and concentration, where applicable, is described in **Supplementary Table 2**.

With such complexity and variety of the factors that can possibly affect the differentiation protocol outcome, we first explore how different protocols affect the composition of the scRNAseq organoids using redundancy analysis (RDA) (**Figure 2C**). We observe the correlation between the protocols described as GSE242057 and GSE142526 and RGC, GSE220624 and Müller Glia, GSE184080 and Cone PhR.

We further discover how timepoint affects the composition of the retinal organoids (**Figure 2D, Figure S5**). As expected, the majority of the samples coming from the similar timepoints cluster together (top right – earlier timepoints, bottom left – later timepoints). While this trend is completely clear for the earlier timepoints, we see some of the samples (GSE220624, GSE236197) to be clustered together with the samples of different timepoint group. As expected, earlier timepoints are characterized by presence of RPC, AC/HC/RGC fate populations, while later timepoints composition can be described with higher frequency of Bipolar/PhR fate, Müller Glia (**Figure S5B**). Noticeably, apatride cells are scattered across the timeline, while lens and non-retinal epithelium are present in early timepoints. On the other hand, such non-canonical/non-retinal populations as non-retinal glia, chimeric PhR, MSX1+ RPC, and astroglial progenitors, are abundant at later timepoints.

### Correlating the timeline of in vitro and in vivo retinal development

We utilize our previously described human fetal retina atlas (HFR)[36] to perform the correlation between in vitro and in vivo timepoints, in both cell class and protocol dependent manner. Human fetal retina atlas contains 108,838 cells from fetal weeks 8 to 27 with a one-week sample resolution with major retinal cell classes present that includes RGC, Bipolar, AC, HC, rod and cone photoreceptors, Müller glia, and RPC.

First, we perform ssGSEA pathway analysis independently on both HROCA and HFR (**Figure 2E**). We observe different trends for necrosis and apoptosis: in HFR these pathways are not enriched in early development but start rising closer to Week 27, while in HROCA both processes initiate from the very beginning (Day 10) of organoid development.

On the other hand, we observe that retinal organoids lose their developmental potential much faster compared to normal retinal development in vivo. For the pathways of optic nerve morphogenesis, neuron fate commitment, and stem cell proliferation, HROCA demonstrates a rapid decrease at Day 20-30, while these pathways remain upregulated in human fetal retina development, where they decrease at Weeks 19-20.

To correlate the timeline of development between in vivo and in vitro, we perform timepoint transfer from HFR to HROCA using scPoli for the retinal populations that are present in vivo development (**Figure 2G**). We demonstrate the predicted timepoints between the cell classes of HFR and cell classes of HROCA at every timepoint group (**Figure 2H**). We observe that:

1. direct correlation of timeline between in vivo and in vitro cannot be applied, i.e. week 1 in vitro is not week 1 in vivo.
2. in vivo and in vitro systems operate with a different concept of time: the trend observed is not completely linear, and while some timepoints in vitro catch up faster to in vivo development, some slack behind. For example, Day 10 to 30 in HROCA is equal to half a week change in HFR, while Day 70 to 90 in HROCA means 2-3 weeks in HFR.
3. in vitro system cannot finish its development: after Day 150 in HROCA there is no positive trend between HROCA and HFR timepoints.
4. cell classes develop asynchronically: while some of the cell classes resemble the general picture of correlated timepoints, RGC in vitro are observed to be lagging behind the general timepoint prediction.

Upon splitting HROCA by protocols, we notice that while some protocols demonstrate similar predicted HFR timepoints for the same HROCA timepoint, there are a few unexpected observations: for Day 60 in HROCA, GSE142526 and GSE235858 match to Weeks 11-12 of HFR, but GSE242057 matches to Week 17. Similarly, at Day 150 HROCA GSE235585 matches to HFR Week 16.5, where GSE184080 is predicted to Week 23 (**Figure S6**).

We also explore transcriptional similarity between the compared identities using Jaccard similarity index (**Figure 2I**) and notice that most of the cell classes in HROCA share the majority of DEGs with HFR. Interestingly, non-retinal or non-canonical cell classes are poorly predicted on HFR, due to these cell classes being absent in vivo development. Among the studies cell classes, the most conservative, or similar, between HROCA and HFR, are cycling RPC (index = 0.32), RPC (index = 0.27), RGC (index = 0.27), rod precursor (index = 0.31), and rod PhR (index = 0.27) cells. While HROCA non-retinal cell classes show minimal similarity to HFR retinal cell classes, we could not identify any profile for apatride cells, unique for HROCA.

The panels of Jaccard similarity index DEG number fine-tuning are available in **Figure S7**.

## Discussion

### The assembly of human organoid atlases

Human stem-cell derived organoids allow to efficiently recapitulate development of multiple tissues and organs, including brain, pancreatic islets, muscle, intestine and others. While the protocols are rapidly developing and vary from lab to lab, the derived organoids are surprisingly consistent, probably due to the fact that the majority of protocols are aimed at the recapitulation of normal development and rely on the cell-cell interactions within these three-dimensional constructs. The single-cell multiomics datasets provided an additional insight at the heterogeneity of the organoids. The relatively high cost per cell of these approaches has been addressed by major undertaking of Atlas assembly, such as human neural organoid cell atlas. [37]

### HROCA composition

Current atlas demonstrates the complexity of existing nature restricted fates within the wild-type retinal organoids differentiated from embryonic and induced pluripotent stem cells. This comprises multiple non-retinal populations, chimeric photoreceptors, immune, lens, and RPE cells. Such composition may be described by the nature of the organoids: their iPSC/hESC origin, potential protocol misguidance, or random perturbations, both cell-intrinsic and -extrinsic.

One of the intriguing populations discovered is a chimeric photoreceptors population, located between photoreceptors and Müller glia on the UMAP as a “bridge” structure. This population has photoreceptor transcriptional pattern (OTX2+ RCVRN+ CRX+ NEUROD1+) but cannot be described as conservative rod or cone photoreceptors, as it expressed markers of both rods (NRL[38,39]) and cones (PDE6H[40], ARR3[41]). Interestingly, this population is also described by CLU[42,43], classical glial marker.

Such population of chimeric photoreceptors that we observe in vitro but not in vivo developing retina system, aligns well with the existing Müller Glia to neurons reprogramming paradigm[44–48] and demonstrates the potential, ability, and sometimes the actual presence of such phenomenon in retinal organoids.

### Batch and scVI-scIB-scANVI fine tuning

Efforts on the large-scale public data integration often face the challenges related to the technological changes spanning throughout the years of sequencing. These can result in platform and chemistry variability, library preparation protocols differences, new versions of reference genome, different depth and coverage of sequencing. Not only that, but also unique experimental design (sorting, enrichment) affects the final dataset composition and its characteristics.

Thus, batch effect on the large-scale atlases becomes the problem amplified to the level where classical batch correction methods [49–54] demonstrate poor integration. Deeper batch correction is required, and this became possible with the application of machine learning methods in computational biology.

We utilized one of these methods, scVI, together with scIB, to perform hyperparameter fine-tuning for better batch integration. scVI offers multiple options for fine-tuning: we applied 4 different settings for n_layers, 3 for n_hidden, 4 for n_latent, 2 for lr. This results in 96 possible combinations. With GPU support, we achieve the results in less than one day, however, all the combinations should be carefully evaluated to select the top performing model.

One could proceed all the way through classical approach up to dimensionality reduction and annotation to identify whether this or another model is more reliable, yet such approach is enormously time consuming. We decided to inspect model validation loss to select top 5 models. In our case, only 2 of them were in the bottom 5% by validation loss, and 3 out of 5 had the n_hidden complexity = 256. We focused on these three models to further inspect more “biology related” metrics provided through scIB, such as batch integration and bio conservation. Through this funnel, we found the top performing scVI model that was further used in downstream analysis.

Such framework performs well and has the advantage of direct, quantitative metrics to investigate, when it comes to atlas integration. However, we would like to notice that every model has its own benefits and gaps. During the analysis, we found that it may be a better way to explore multiple models at the same time, as some populations separate better or worse in different models, and to merge the annotation from multiple models prior to scANVI analysis.

### Protocols metadata

A primary objective of HROCA is to characterize the diversity of retinal organoid differentiation protocols reported across the field. While published literature remains the principal source of such protocols, extracting comprehensive and standardized metadata presents notable challenges. Although many groups have made commendable efforts to enhance transparency and reproducibility by including detailed protocol descriptions, inconsistencies remain. In several cases, crucial metadata are missing, timelines of differentiation vary considerably between studies, and nomenclature for key reagents is not standardized.

These discrepancies introduce subjectivity and hinder large-scale protocol comparisons. To mitigate this, our approach emphasized preserving the original logic and terminology used by the authors rather than reinterpreting or transforming the data into a harmonized framework. While this strategy prioritizes fidelity to the source material, it also results in unavoidable data loss—particularly when ambiguous or insufficiently defined elements were labeled as “NA” to avoid misclassification. In cases where labels carried potential dual meanings, we refrained from reassigning them to prevent erroneous assumptions. For example, when one protocol referred to a supplement as a “WNT inhibitor” and another specifically listed “SU5402,” we retained the original designations rather than merging them, thus maintaining resolution at the cost of immediate comparability.

Even with detailed protocol documentation, certain critical pieces of information remain inaccessible. One notable example is the tissue origin of iPSCs or hESCs, a factor potentially influencing organoid differentiation outcomes. Furthermore, variation in organoid composition is likely driven by factors extending beyond the differentiation protocols themselves. These may include, but are not limited to, differences in library preparation protocols, organoid selection procedures, cell viability and cDNA concentrations, as well as batch effects arising from assay conditions, sequencing technologies, and institutional infrastructure. These confounding variables underscore the complexity of achieving standardization and highlight the importance of continued efforts to improve protocol reporting and metadata completeness across the field.

### Nature restricted cell classes and states

In its current version, HROCA captures data exclusively from wild-type retinal organoids. This limitation prompted us to explore which cell states naturally emerge and which do not - what we define as “nature restricted” states. An example is the appearance of Chimeric Photoreceptor class in HROCA but not in the HFR Atlas. These cells occupy an intermediate position in the UMAP space between Müller glia and photoreceptors and co-express markers from both lineages: PDE6H (cone photoreceptor), NRL (rod photoreceptor), and CLU (a Müller glia marker). The presence of these hybrid states may suggest a latent reprogramming potential of human retinal cells, at least at the transcriptional level, consistent with prior findings in zebrafish and murine systems[44–48] .

This observation raises a broader question: what are the theoretical boundaries of cellular identity? If we were to imagine a hypothetical cell expressing an implausible combination of unrelated markers occupying an unpopulated region in the UMAP space its absence may signal a “naturally impossible state.” Such a state would likely violate core principles of lineage commitment and cell fate logic, and thus may not exist even under perturbed conditions.

HROCA, therefore, provides insight into both the flexibility and the constraints of retinal cell identities in vitro. While it reveals cell states that may not be observed in vivo, including transitional or hybrid populations, these findings remain grounded in biologically plausible trajectories. Looking forward, the boundaries of these natural limits may shift in HROCA v2, where we intend to incorporate organoids derived from disease models or subjected to experimental perturbations. Furthermore, expanding the platform to include other in vitro systems - such as 2D cultures, retinal explants, co-culture models, and assembloids - could offer additional resolution into how context and environmental complexity influence the emergence or suppression of specific cell states. It may further enhance our understanding by bringing additional conditions intp the system, such as post-transplantation, hypoxic, maldifferentiated organoids into the Atlas.

### Apatride cells

Among the unique cell states identified in HROCA, one particularly intriguing population is what we have termed “apatride cells.” This label—derived from the notion of statelessness—reflects the inability to assign these cells to any known lineage, as they do not express markers characteristic of any canonical retinal cell type. We rigorously evaluated the possibility that these cells result from technical artifacts. However, they display normal transcriptomic quality, including typical numbers of detected transcripts and genes, as well as expected proportions of mitochondrial and ribosomal gene expression.

Importantly, these cells do not show elevated expression of genes associated with cell death, inflammation, or hypoxic stress, further supporting their biological rather than artifactual nature. As such, we hypothesize that apatride cells may represent a “fateless” or “doomed” cell state—cells that have not yet, or may never, commit to a defined lineage. While this interpretation remains speculative, their consistent presence across multiple timepoints and increasing abundance at later stages suggest that their emergence is not random. Rather, it may reflect intrinsic limitations or inefficiencies within the organoid differentiation process.

Understanding the origin and potential trajectories of these cells will require further investigation. In particular, lineage tracing experiments could clarify whether apatride cells are truly developmentally arrested or whether they represent transient, pre-committed states that ultimately adopt a defined identity under appropriate conditions.

Their existence underscores the complexity and heterogeneity of in vitro retinal development and highlights the need to better understand noncanonical or intermediate states that may not be captured in standard annotations.

## Funding

Gilbert Family Foundation–GFF00 (P.B.), Department of Defense - VRP FTTSA (P.B.), MGB GCTI Grant (P.B.)

## Data availability statement

The software and packages used in this study are available in Supplementary Table 5. The complete code, trained models, and multiple .h5ad/.h5Seurat objects will be available on GitHub upon publication. CELLxGENE deposited HROCA will be available upon publication.

## Author contributions

E.K., E.L., N.R., P.B. performed the research; E.K., E.L. analyzed the data and created the figures; E.L., E.K. performed public data parsing and reanalysis; E.L., N.R., E.K. performed the protocol-related part of the study; E.K., P.B. wrote the paper; and P.B. supervised the research.

## Competing interests

The authors declare no conflict of interest.

---

**Figure S1.**
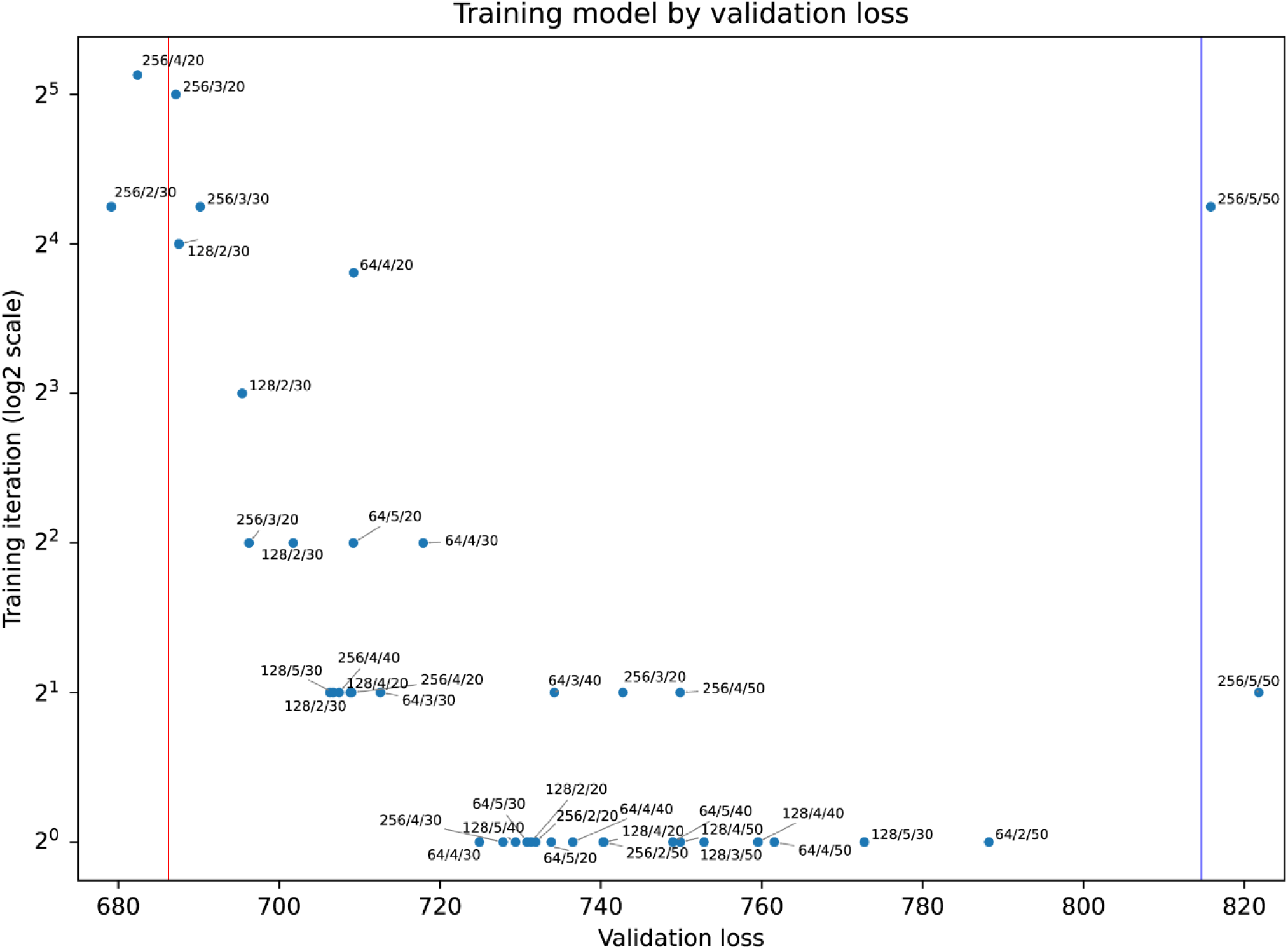
Scatterplot demonstrating validation loss for every trained scVI model.

**Figure S2.**
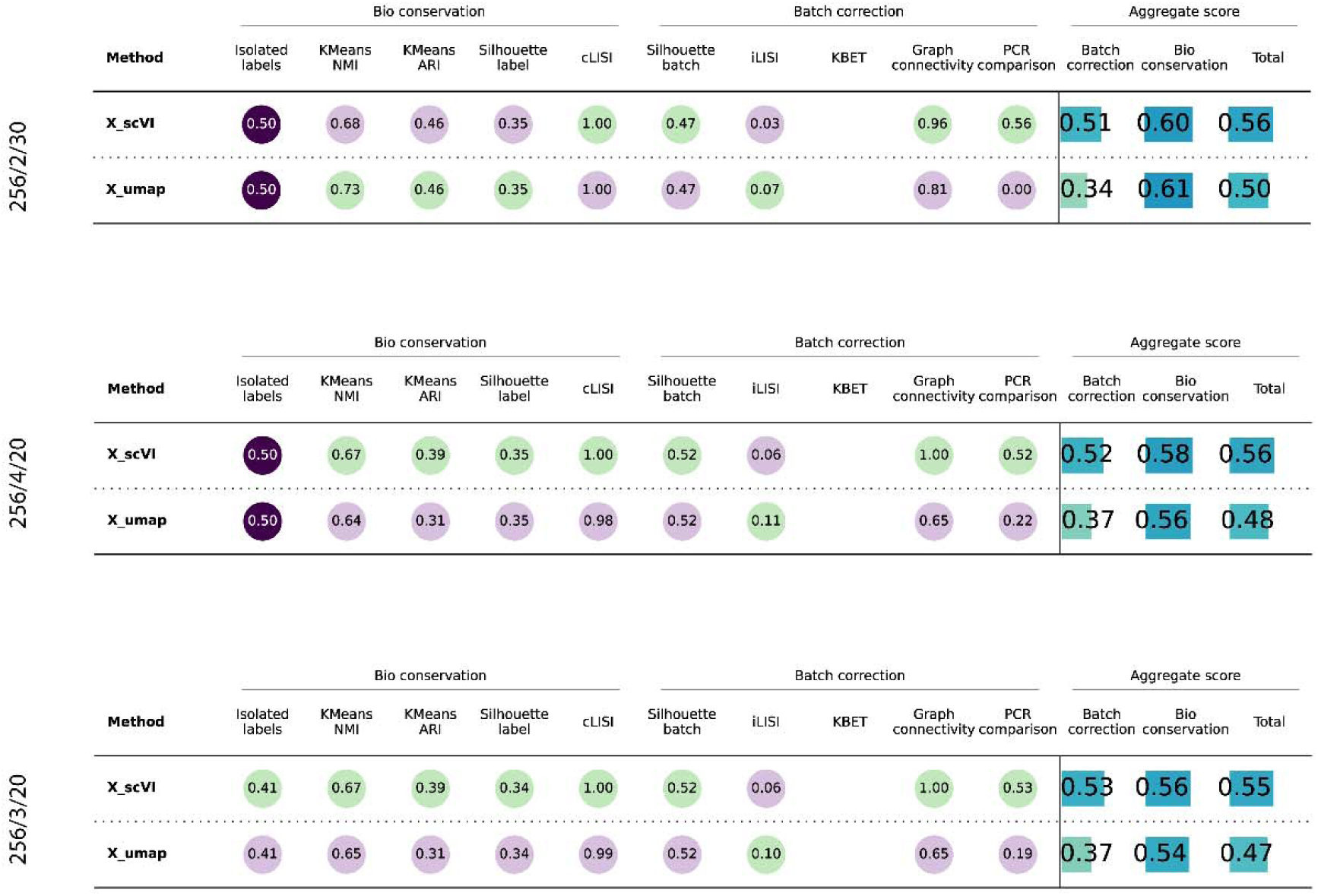
Benchmarking parameters for the top 3 scVI models.

**Figure S3.**
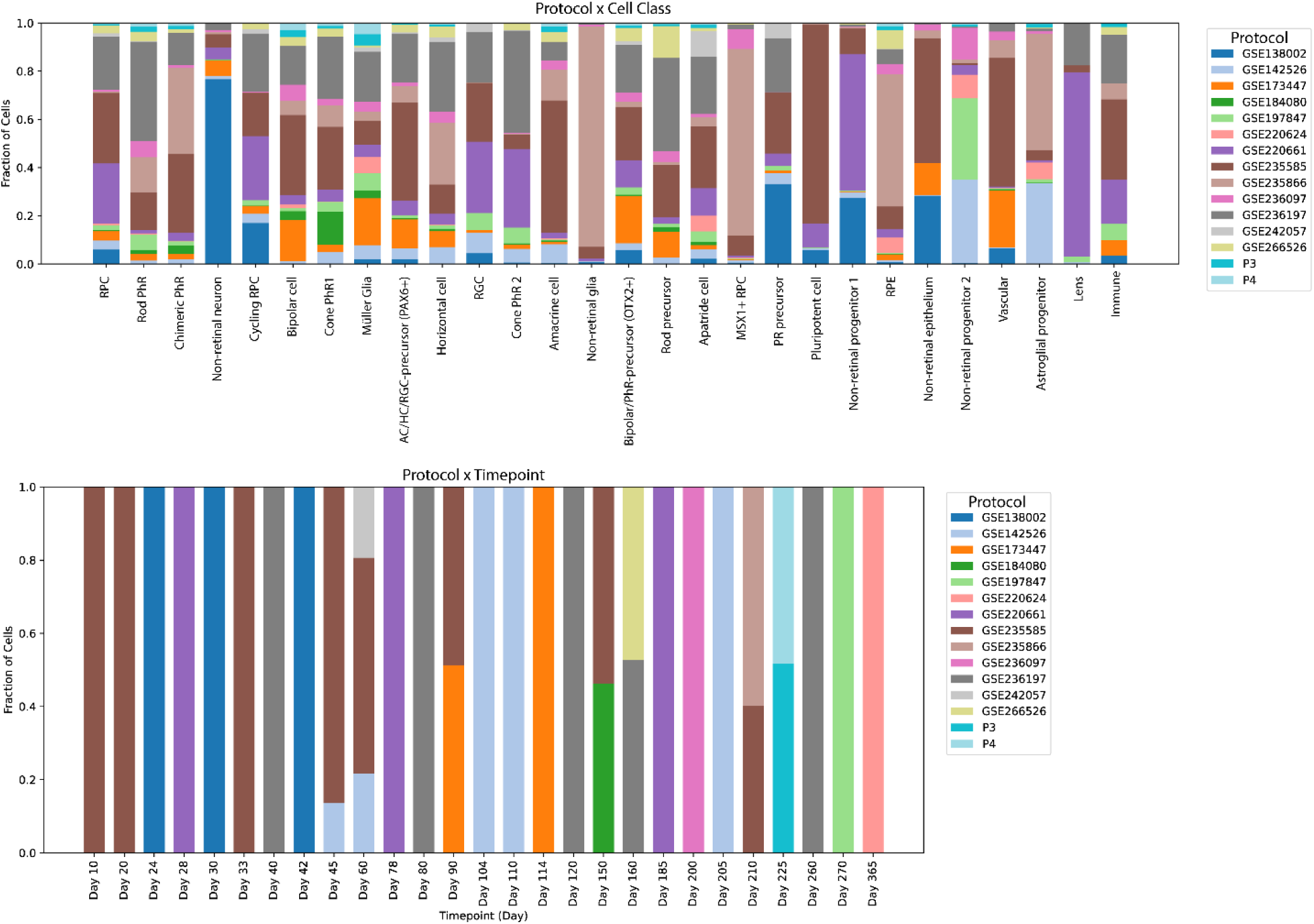
Bar charts demonstrating the distribution of protocols across cell classes and timepoints.

**Figure S4.**
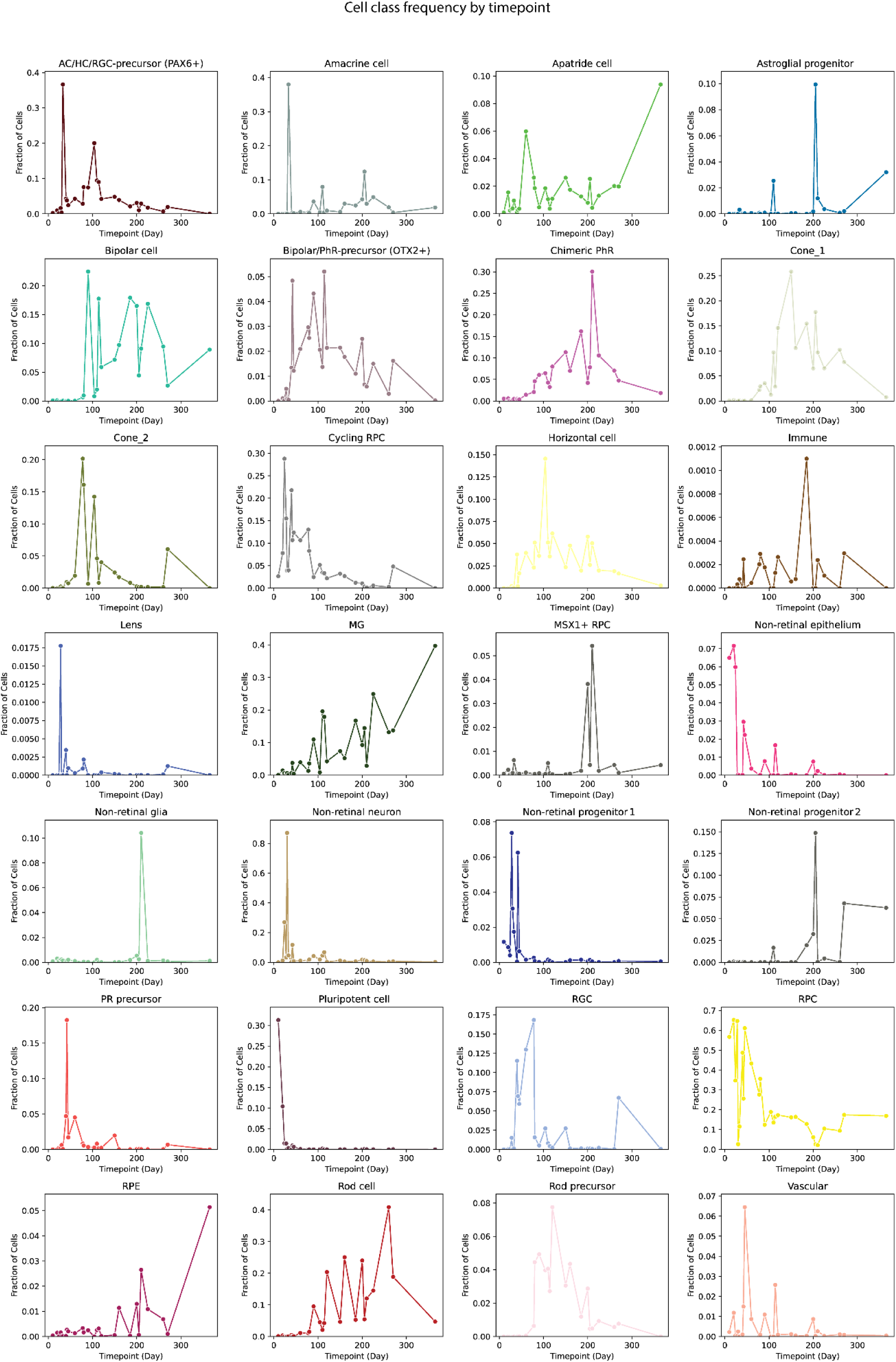
Charts representing the frequency of every HROCA cell class across the timeline from day 10 to day 365.

**Figure S5.**
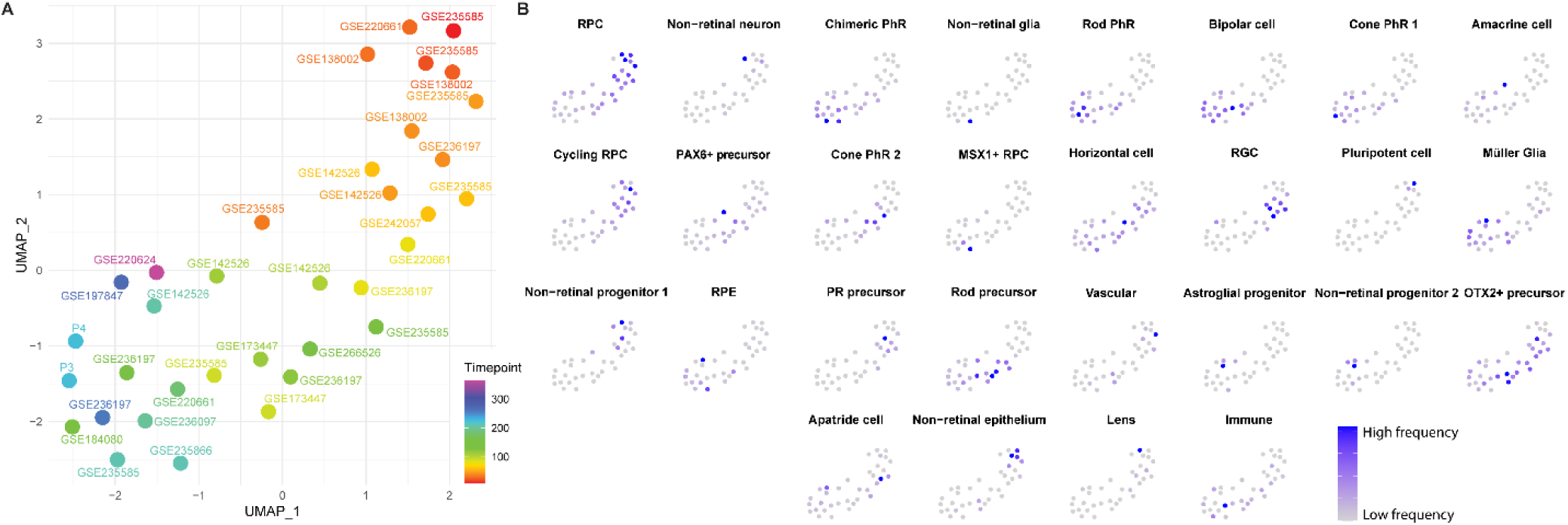
HROCA protocols and timepoints analysis uncovers the pattern of cell classes arising. (A) UMAP embedding of protocol compositions. Each dot represents a sample that belongs to a protocol and a timepoint. Color codes the timepoint. (B) Feature plots for every cell class on the embedding. Color codes the frequency of the cell class.

**Figure S6.**
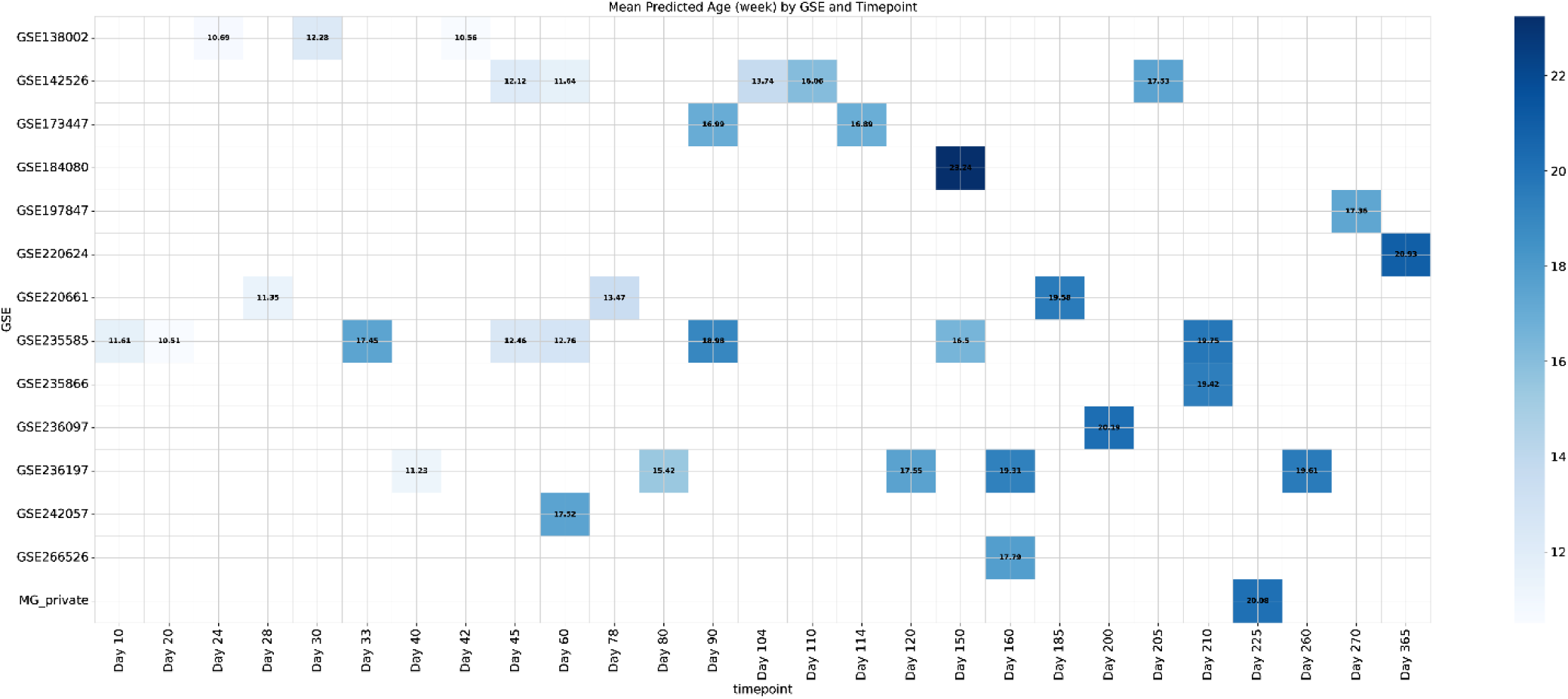
scPoli-based prediction of HROCA to HFR timepoints split by individual protocols and timepoints.

**Figure S7.**
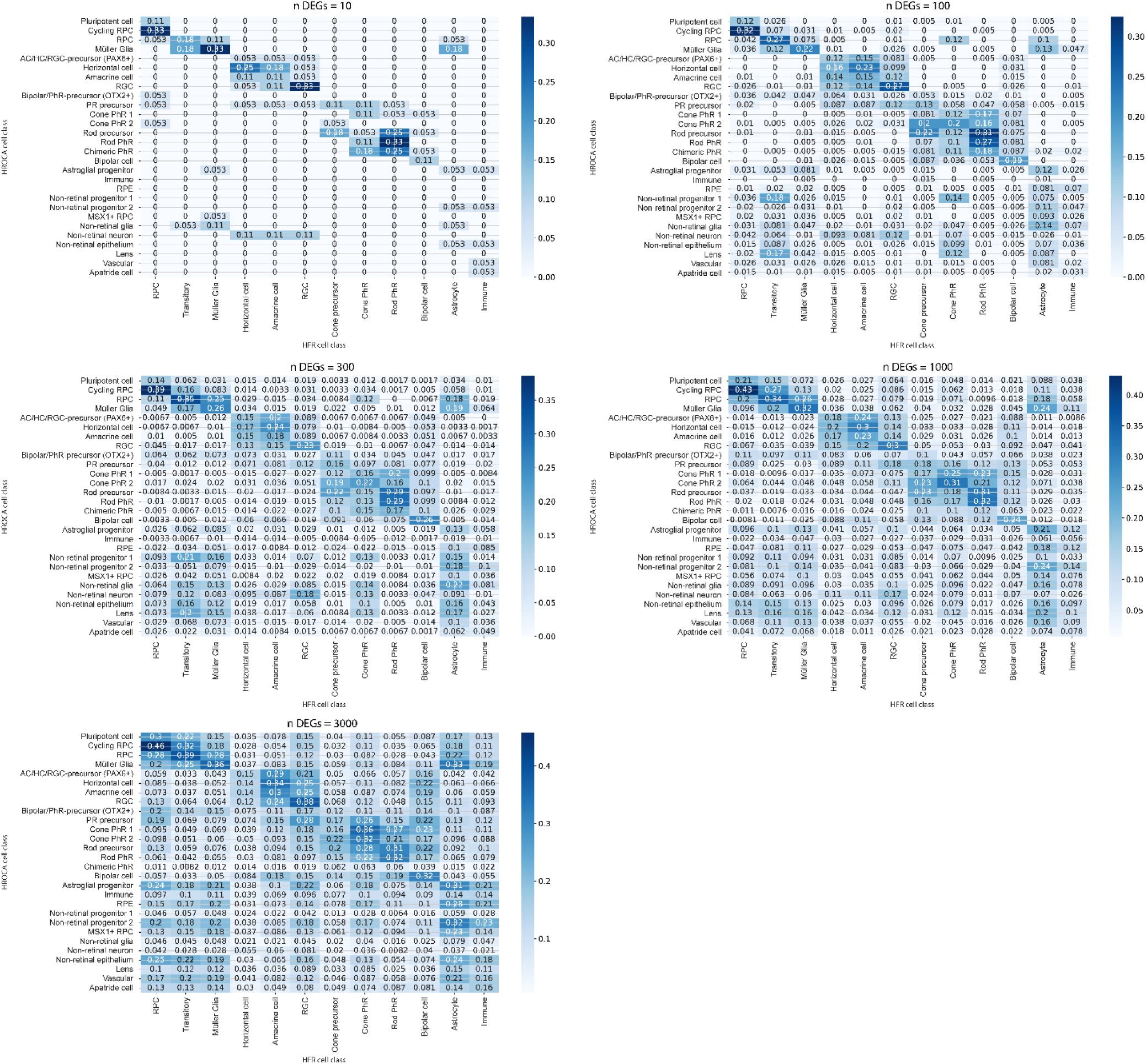
Jaccard similarity index heatmap generated for 10, 100, 300, 1000, and 3000 DEGs per cell class.

## Notes

### Competing Interest Statement

The authors have declared no competing interest.

