## Supplementary figures and images for "Human retinal organoid single-cell atlas allows to reconstruct retinal development at high resolution and identify nature restricted transcriptional states in vitro"

### Figure S1

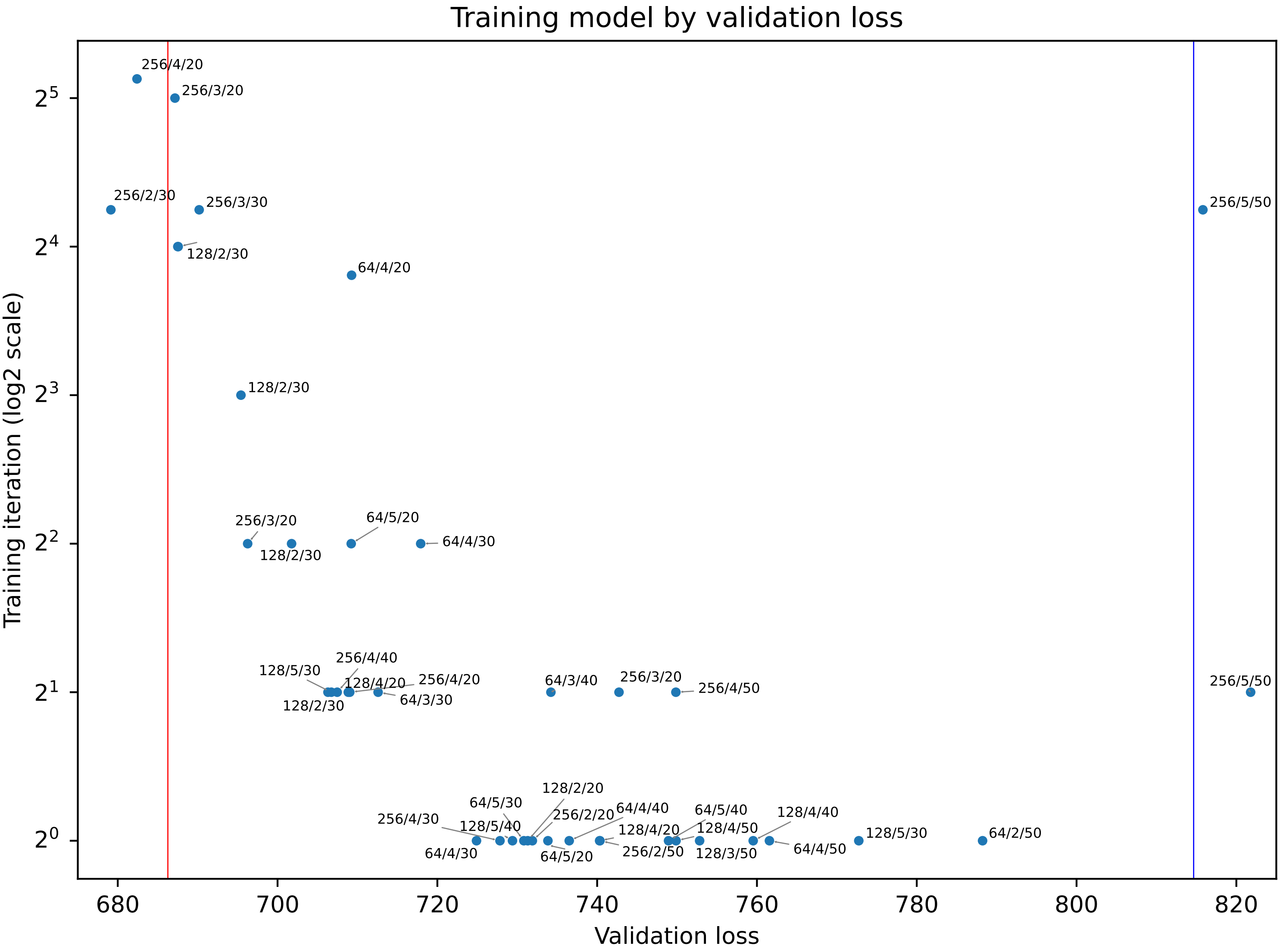

### Figure S3

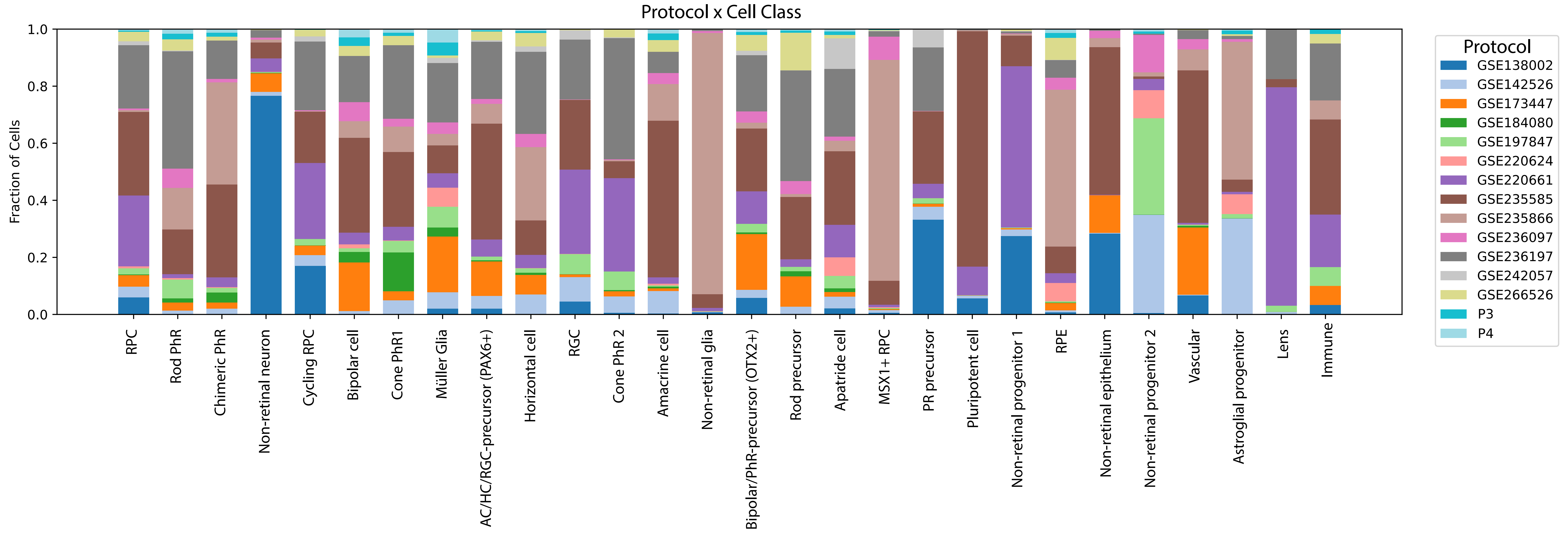

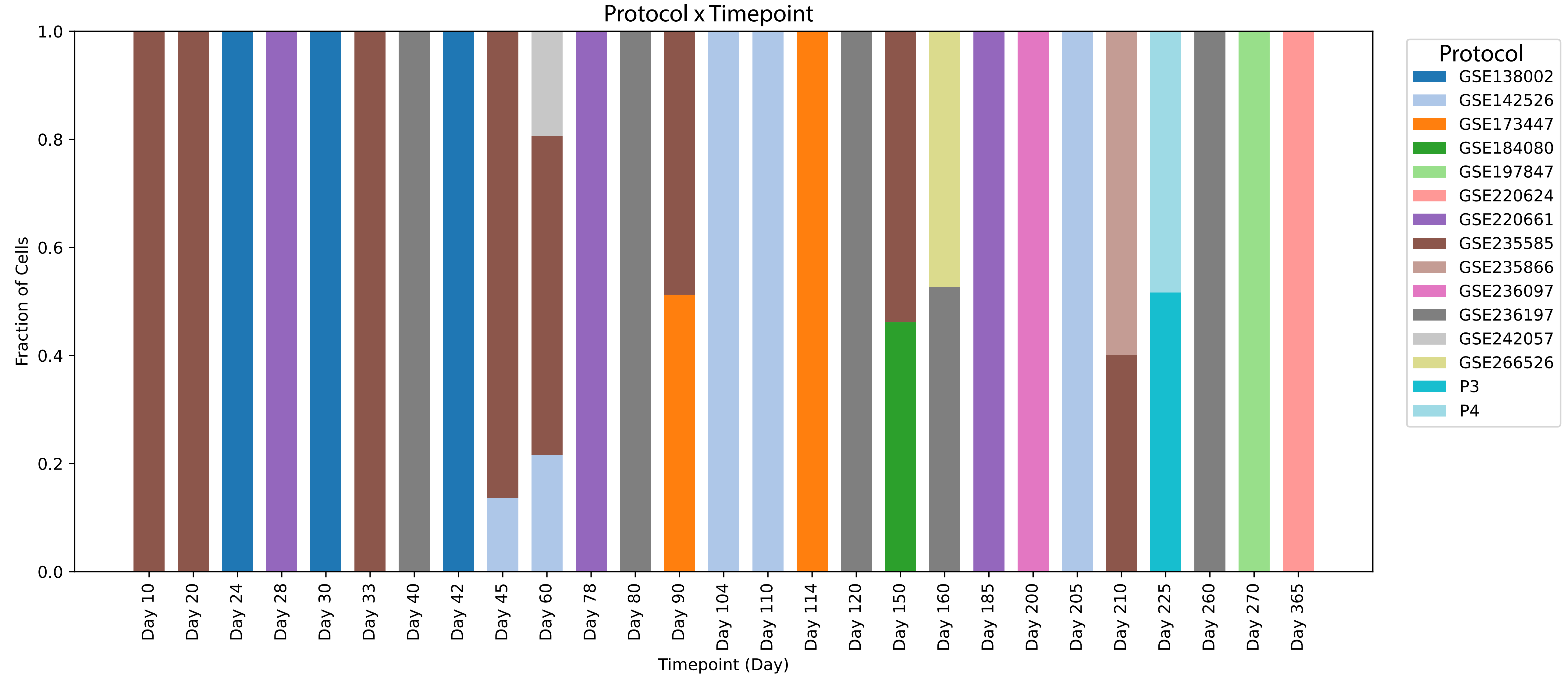

### Figure S4

#### Cell class frequency by timepoint

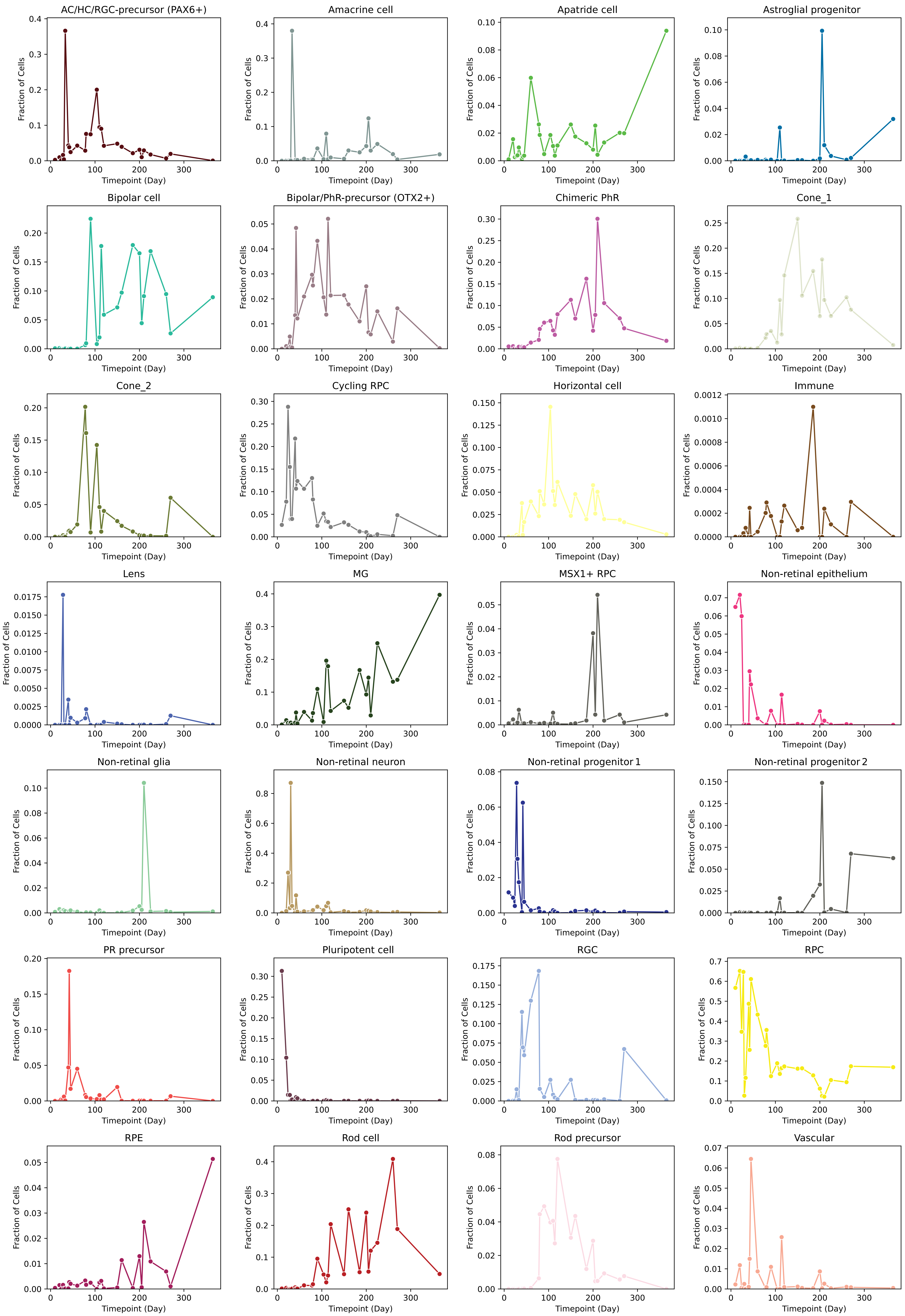

### Figure S5

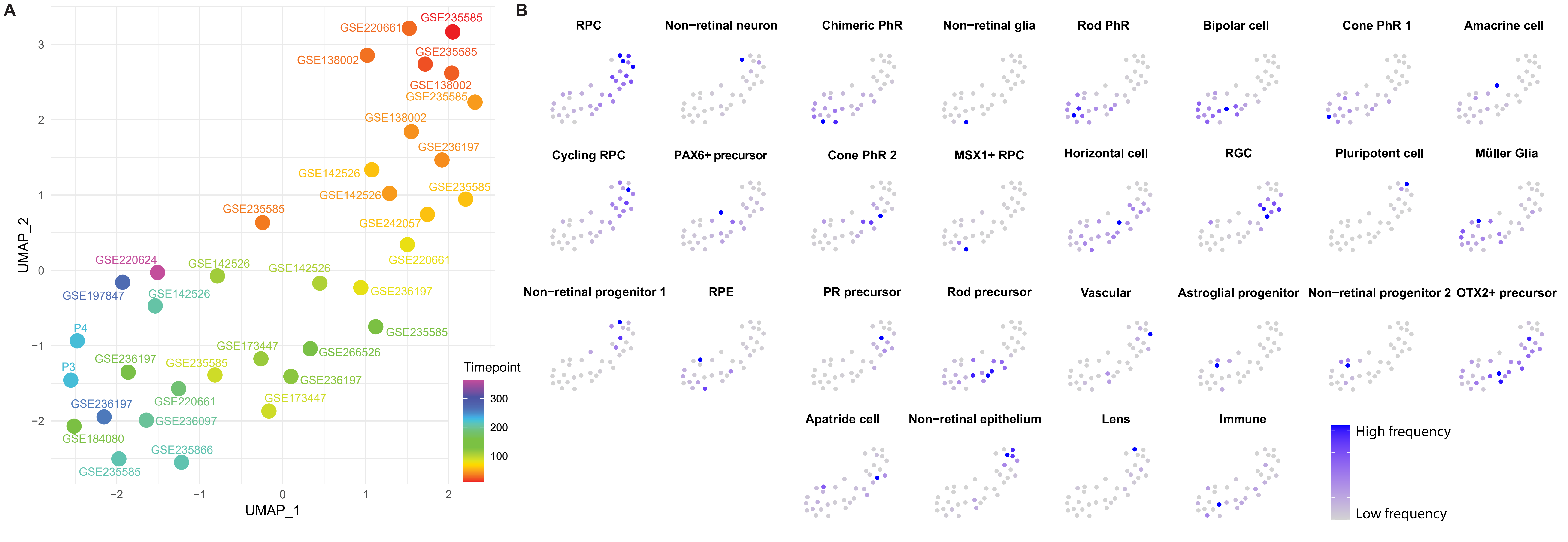

### Figure S6

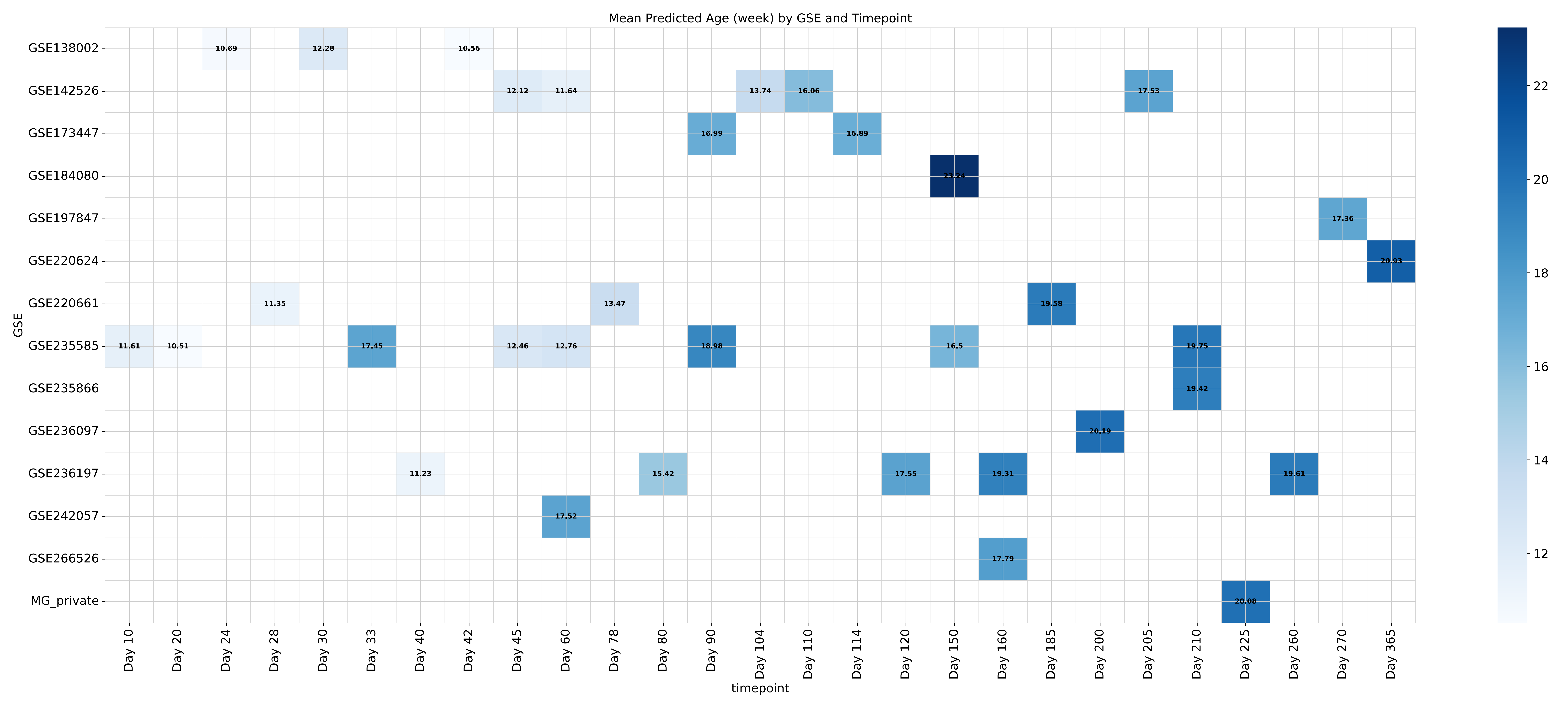

### Figure S7

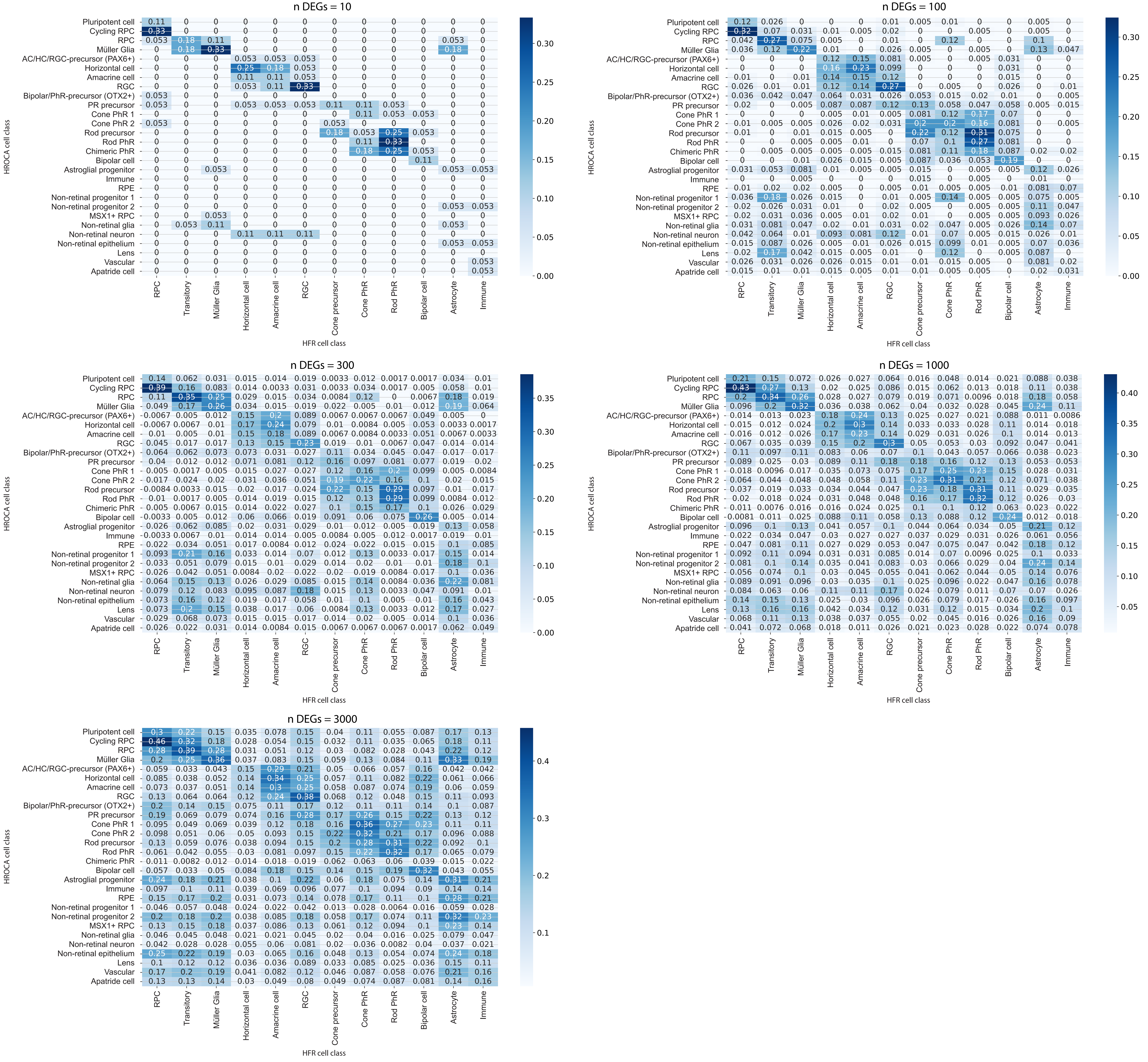
