## Supplementary material for "Human retinal organoid single-cell atlas allows to reconstruct retinal development at high resolution and identify nature restricted transcriptional states in vitro": Figure S2

256/2/30

| Method | Bio conservation | | | | | Batch correction | | | | | Aggregate score | | |
|---|---|---|---|---|---|---|---|---|---|---|---|---|---|
| | Isolated labels | KMeans NMI | KMeans ARI | Silhouette label | cLISI | Silhouette batch | iLISI | KBET | Graph connectivity | PCR comparison | Batch correction | Bio conservation | Total |
| X_scVI | 0.50 | 0.68 | 0.46 | 0.35 | 1.00 | 0.47 | 0.03 | | 0.96 | 0.56 | 0.51 | 0.60 | 0.56 |
| X_umap | 0.50 | 0.73 | 0.46 | 0.35 | 1.00 | 0.47 | 0.07 | | 0.81 | 0.00 | 0.34 | 0.61 | 0.50 |

256/4/20

| Method | Bio conservation | | | | | Batch correction | | | | | Aggregate score | | |
|---|---|---|---|---|---|---|---|---|---|---|---|---|---|
| | Isolated labels | KMeans NMI | KMeans ARI | Silhouette label | cLISI | Silhouette batch | iLISI | KBET | Graph connectivity | PCR comparison | Batch correction | Bio conservation | Total |
| X_scVI | 0.50 | 0.67 | 0.39 | 0.35 | 1.00 | 0.52 | 0.06 | | 1.00 | 0.52 | 0.52 | 0.58 | 0.56 |
| X_umap | 0.50 | 0.64 | 0.31 | 0.35 | 0.98 | 0.52 | 0.11 | | 0.65 | 0.22 | 0.37 | 0.56 | 0.48 |

256/3/20

| Method | Bio conservation | | | | | Batch correction | | | | | Aggregate score | | |
|---|---|---|---|---|---|---|---|---|---|---|---|---|---|
| | Isolated labels | KMeans NMI | KMeans ARI | Silhouette label | cLISI | Silhouette batch | iLISI | KBET | Graph connectivity | PCR comparison | Batch correction | Bio conservation | Total |
| X_scVI | 0.41 | 0.67 | 0.39 | 0.34 | 1.00 | 0.52 | 0.06 | | 1.00 | 0.53 | 0.53 | 0.56 | 0.55 |
| X_umap | 0.41 | 0.65 | 0.31 | 0.34 | 0.99 | 0.52 | 0.10 | | 0.65 | 0.19 | 0.37 | 0.54 | 0.47 |
